# Identification and characterization of a Fibrillin-1 derived matrikine for cardiac regeneration and repair

**DOI:** 10.1101/2025.11.27.690884

**Authors:** Kyle J. Edmunds, Elizabeth C. Porter, Yan-Ru Lai, Jacques Guyette, Corin Williams, Ajith Jaiganesh, Harald Ott, Justin S. Weinbaum, Lauren D Black

## Abstract

The development of regenerative strategies to repair the heart is of high importance. Our lab has shown that extracellular matrix derived from decellularized fetal myocardium promotes neonatal cardiomyocyte proliferation in vitro. The goal of this study was to identify specific peptide(s)/protein(s) in solubilized cardiac ECM responsible for this proliferative effect. We hypothesized that isolation and then treatment with one or more small synthetic peptide derived from this source could replicate the cellular response to whole solubilized ECM. Decellularized fetal and adult rat hearts were fractionated by molecular weight using SDS-PAGE and transferred to PVDF membranes. Analysis of cardiomyocytes cultured on the membranes revealed regions of enhanced cardiomyocyte proliferation. Subsequent isolation and proteomic analysis of the protein bands that that correlated with proliferative regions identified fibrillin-1 as the predominant ECM protein associated with these regions of cardiomyocyte proliferation. One region (residues 55-86) of fibrillin-1 was synthesized as a peptide and tested for a direct effect on cardiomyocyte proliferation. Compared to positive and negative controls, as well as scrambled and alkylated versions, this peptide led to 3-4 fold increase in cardiomyocyte proliferation. Analysis of the amino acid sequence demonstrated high homology with laent-TGF-β binding proteins and subsequent experiments showed that the matrikine could also reduce TGF-β induced activation of cardiac fibroblasts. These data suggest that individual peptides derived from soluble ECM could have utility as a novel therapeutic for cardiac tissue engineering and regeneration.

## INTRODUCTION

The extracellular matrix (ECM) is increasingly recognized as a dynamic signaling hub rather than a passive structural scaffold. Beyond providing mechanical support, the ECM orchestrates cellular behaviors through biochemical cues (1), mechanotransduction (2), and the release of bioactive fragments known as matrikines (3). These cryptic peptides, generated during proteolytic remodeling, have emerged as potent regulators of processes such as angiogenesis (4), immune modulation (5), and tissue repair (6). In development and disease, ECM-derived signals integrate with growth factor pathways and metabolic programs to shape cell fate decisions, yet their contribution to regenerative responses remains incompletely understood.

Cardiac regeneration exemplifies this complexity. Unlike lower vertebrates, the adult mammalian heart exhibits minimal capacity for functional repair following injury (7), largely due to the limited proliferative potential of postnatal cardiomyocytes (8). However, fetal and neonatal hearts retain a transient window of regenerative competence (9), suggesting that developmental cues—including those embedded within the cardiac ECM (10)—could be harnessed therapeutically. Indeed, ECM components such as agrin (11), periostin (12), and neuregulin (13) have been implicated in cardiomyocyte proliferation and maturation, while emerging evidence points to matrikines as additional modulators of myocardial repair (14).

ECM signaling in regeneration can be envisioned as a multi-tiered process. At the structural level, intact ECM provides mechanical cues that influence cell shape and cytoskeletal tension (15). At the molecular level, full-length ECM proteins engage receptors such as integrins and syndecans to activate intracellular pathways. During tissue remodeling—whether developmental or injury-induced—proteolytic cleavage generates matrikines, which function as soluble effectors capable of diffusing through the microenvironment and modulating cell behavior. These fragments can mimic growth factor activity, alter metabolic states, and fine-tune immune responses (3), creating a localized niche conducive to repair. Moreover, the mechanism of action of matrikines may be complex as evidence suggests that, beyond mimicking growth factors, they may also act through integrin-mediated signaling (16), metabolic reprogramming (17), or canonical pathways such as Wnt/β-catenin (16). Understanding this hierarchy—intact ECM, structural signaling, and matrikine-mediated biochemical cues—offers a roadmap for designing biomaterials and therapeutics that recapitulate regenerative microenvironments.

Despite these insights, most studies interrogate individual ECM proteins, overlooking the combinatorial and context-dependent nature of matrix signaling. Perfusion-decellularized scaffolds (18) and solubilized ECM preparations (19) offer a more holistic approach, preserving native architecture and composition. Using such strategies, fetal and neonatal cardiac ECM has been shown to enhance cardiomyocyte proliferation – both in insoluble (20) and soluble (10) forms - and improve post-infarct remodeling compared to adult ECM (21), implicating developmental stage–specific bioactivity. Notably, partial proteolysis of adult ECM can restore proliferative effects (22), raising the possibility that matrikines generated during digestion are key effectors.

Here, we present a platform for systematic discovery of ECM-derived peptides with regenerative potential. By integrating SDS-PAGE fractionation, direct cardiomyocyte culture on protein blots, and LC-MS/MS analysis, we identify candidate fragments that modulate cell phenotype. Applying this approach to cardiac ECM, we uncovered a fibrillin-1–derived peptide that significantly promotes cardiomyocyte proliferation. These findings highlight matrikines as a previously underexplored class of bioactive molecules and suggest new avenues for leveraging ECM signaling in cardiac repair and beyond.

## RESULTS

### Optimization of cECM Peptide Separation and Blot Culture

To identify cECM peptides that induce specific cellular phenotypes, we first developed a novel method to culture primary cardiac cells directly on a protein blot of fractionated cECM peptides. To accomplish this, fetal and adult rat hearts were isolated and decellularized using developmental age specific procedures previously confirmed to remove DNA and other cellular components while retaining most of the native cECM (10, 18, 19). Once the tissue was devoid of cellular material, cECM was digested into peptides via solubilization with pepsin (19). Next, solubilized cECM peptides were fractionated by molecular weight using standard sodium dodecyl sulphate–polyacrylamide gel electrophoresis (SDS-PAGE) methods for cellular protein separation Briefly, the cECM was sonicated, treated with dithiothreitol (DTT) and loaded onto the gel. Samples were intentionally loaded as mirror images on the left and right halves of the gel, so as to control for differential fractionation as a result of variations in electrophoresis time. Initial experimentation suggested that the maximum loading capacity of cECM in each well that still allowed for efficient separation of the peptides was 450μg of cECM per lane (**Supplemental Figure 1**). After running the PAGE, half of the resulting gel was transferred to a polyvinyl difluoride (PVDF) membrane. The second half of the gel was kept for extraction of peptides and subsequent LC-MS/MS compositional analysis. Finally, the transferred membrane was sterilized via penicillin-streptomycin rinses and then blocked with bovine serum albumin (BSA) before being placed in culture medium. Note that the removal of DTT from the solution allows the peptides to reform disulfide bonds (23). Primary neonatal rat cardiac cells were isolated as previously described (24), seeded, and cultured directly on the surface of the membrane containing the transferred fractionated cECM.

### Detection of Proliferative CM Regions within the Blot Culture

To identify molecular weight regions, or “hotspots,” potentially containing peptides that induce a proliferative phenotype in neonatal rat cardiomyocytes, the cardiac cell blot culture was fixed, permeabilized, and blocked after either one or five days in culture. The cells were immunostained for the proliferation marker Ki67 and cardiac α-actinin to co-label proliferative cells and CMs, respectively. The fixed blot culture was then imaged using an inverted fluorescence microscope. After acquisition, the membrane was reconstructed by alignment of sequential images. We observed cell attachment for the full range of molecular weight on the membrane (**Figure 1A**). Furthermore, subsequent analysis of the images with a custom CellProfiler pipeline determined the number of cells (**Figure 1B**) and the number of CMs (**Figure 1C**) as a function of molecular weight. The total numbers of cell nuclei, Ki-67, and cardiac α-actinin positive cells in each lane and as a function of molecular weight were determined using customized pipelines in CellProfiler. Averaging the CM density across the different lanes of the same developmental aged cECM demonstrated that, while variable, there were no significant differences in density dependent on molecular weight (**Figure 2**). In addition, the overall average CM density for the entire blot (dashed line), for both ECM ages, and across two different experiments was similar for both fetal cECM (**Figure 2A, B**) and adult cECM (**Figure 2C, D**). By concentrating on the top 5% of the distribution of proliferative CM number (**Supplemental Figure 2**), we identified defined molecular weight regions (black bars) that corresponded to large numbers of proliferative CMs in both fetal cECM (**Figure 2E, F**) and adult cECM (**Figure 2G, H**) preparations. Locations of these regions also correlated with higher overall numbers of CMs (**Figure 2A-D**).

**Figure 1:**
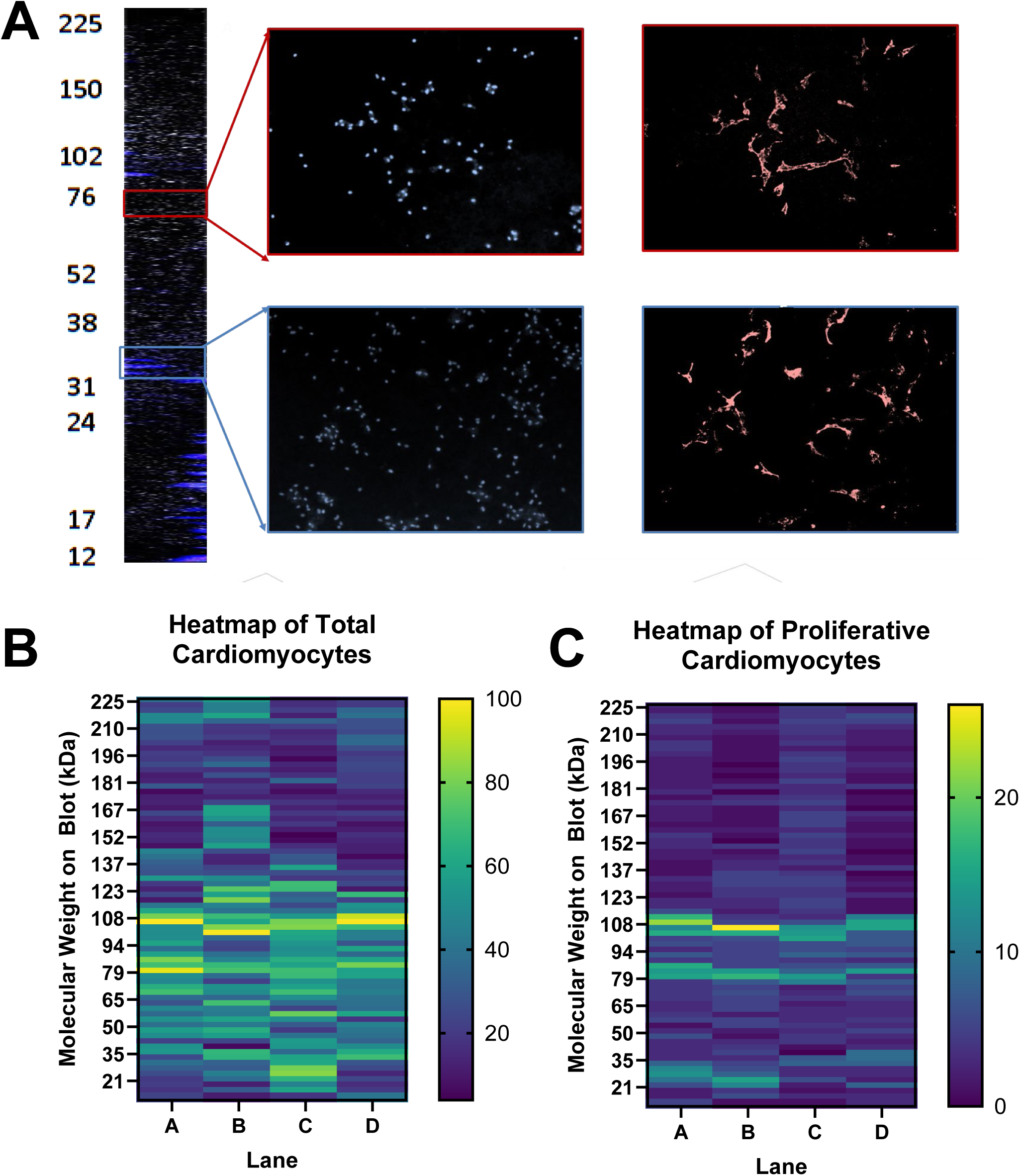
Neonatal Cardiac Cells can be cultured on fractionated cardiac extracellular matrix blots. **(A)** A composite of 77 stitched images demonstrates significant cell adhesion along the entire lane from high molecular weight (225 kDa) to low molecular weight (12 kDa) on the blot of fractionated cardiac extracellular matrix. Zooming in we can identify regions with lower amounts of cell attachment (red boxes, top right) and other areas with higher amounts of cell attachment (blue boxes, lower right). Note that cardiomyocytes are labeled with cardiac α-actinin (red) and nuclei are labeled with DAPI (blue). When 4 lanes of the same ECM are run on the same blot (lanes A-D) we see that while there is some variation lane-to-lane, generally both total cardiomyocyte number **(B)** and proliferative cardiomyocyte number **(C)** are relatively consistent.

**Figure 2:**
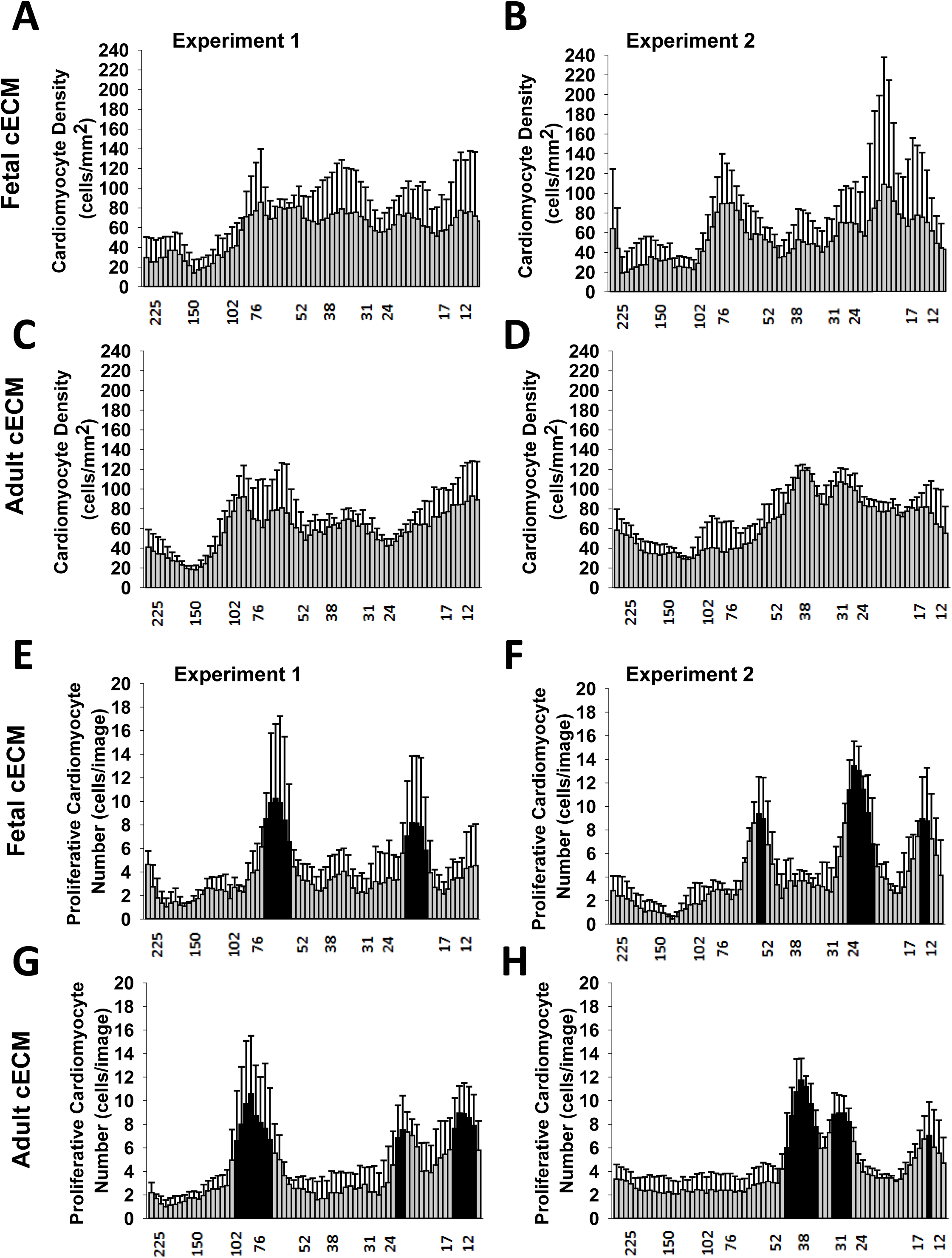
Quantification of blot cultures for fetal and adult cardiac ECM across multiple experiments demonstrates specific regions of high levels of cardiomyocyte proliferation. Quantification of cardiomyocyte density as a function of molecular weight on the blot for 4 lanes of the PVDF membrane for 2 separate experiments of fetal cardiac extracellular matrix **(A,B)** and adult cardiac extracellular matrix **(C,D)** validates relatively consistent attachment of cells along the molecular weight axis. In addition, the mean value for cardiomyocyte for the length of the blot (dashed line in each plot) was consistent among both ECM developmental ages and experiments. Quantification of proliferative cardiomyocyte density as a function of molecular weight on the blot for 4 lanes of the PVDF membrane for 2 separate experiments of fetal cardiac extracellular matrix **(E,F)** and adult cardiac extracellular matrix **(G,H)** demonstrated specific regions of higher proliferative activity. The bars filled in black represent the top 5% of values of the overall blots’ distribution (see **Supplemental Figure 2**), indicating significantly higher levels of proliferation in these regions. While the specific molecular weight regions are slightly different between experiments and ECM types, the overall degree of proliferation in these regions is similar and the mean value of the blots (dashed line for each plot) are also similar between ECM types and experiments.

### Proteomic Analysis for Identification of cECM Peptides that Promote CM Proliferation

To assess whether a particular peptide or peptides led to the proliferative response of the CMs in our blot culture system, the relevant molecular weight regions identified in in **Figure 2**, were excised from the second half of the electrophoresed gel. The peptides present in these regions were then eluted from the gel, processed, and sent for proteomic analysis via LC-MS/MS. Regions corresponding to comparatively low numbers of proliferative CMs were also excised and analyzed as controls. Resulting spectral data was analyzed in order to quantify changes in relative amounts of various proteins present in the excision regions. The proteomic results showed that the only cECM protein that was significantly higher in both fetal and adult proliferative excisions as compared to control excisions was fibrillin-1 (**Figure 3A**). Both fetal and adult proliferative excisions contained significantly less collagen I α-2 protein than control excisions (**Figure 3A**). Additionally, adult proliferative excisions also contained significantly less collagen I α-1 than either the fetal proliferative excisions or controls (**Figure 3A**). While the reduction of collagen −1 peptides could explain changes in CM proliferation if those peptides were to inhibit proliferation, we hypothesized that the significant increase of peptides from fibrillin-1 was the likely reason for the observed increase in CM proliferation in the specific molecular weight regions.

**Figure 3:**
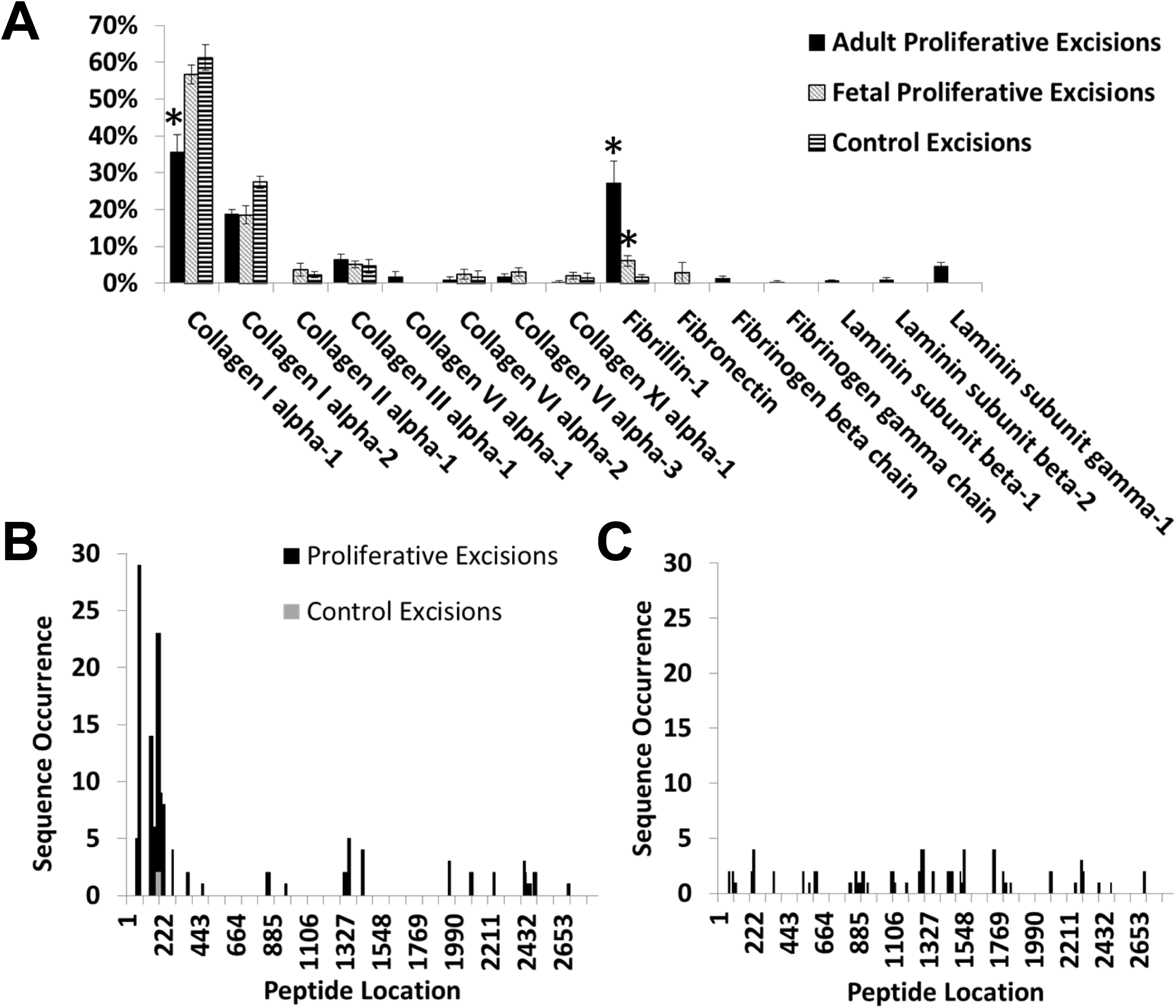
Proteomic analysis identified fibrillin-1 peptides in high abundance in the proliferative regions of the blot culture. **(A)** Percent of ECM protein Spectral counts for various ECM proteins found in adult ECM proliferative regions (Black Bar), fetal ECM proliferative regions (Gray Bar) and control regions (Striped Bar). Note that the only protein that was in higher abundance in both adult and fetal ECM proliferative regions was fibrillin-1. * denotes p<0.05 compared to control excisions. **(B)** Mapping the fibrillin-1 peptide sequences from proliferative and control excisions into the full-length sequence for fibrillin-1 showed a higher abundance of peptides near the N-terminus of the fibrillin-1 protein. Mapping fibrillin-1 sequences identified from whole fetal ECM into the full-length fibrillin-1 sequence showed relative even distribution of peptides throughout the protein, indicating that the peaks found in the proliferative mapping are not due to identification restraints from ProteinPilot.

To further identify the specific sequence of the peptide or peptides from fibrillin-1 that were present in the proliferative regions of the blot culture, we mapped the returned sequences from the LC-MS/MS identification to the known full-length sequence of fibrillin-1 of the Sprague-Dawley rat. The primary returned sequences were clustered near the N-terminus of the fibrillin-1 protein and the largest peaks corresponded to amino acid residues 55 to 232 (**Figure 3B**). Furthermore, proteomics analysis of whole, unfractionated cECM solubilized with urea, gave a relatively homogenous spectrum of fibrillin-1 results (**Figure 3C**), suggesting that the observed peaks near the N-terminus were not an artifact of the software’s ability to identify fibrillin-1. We did observe a small gap in sequence occurrence between the front most of these peaks (55–86) and the others (139–232). Since the fractionated cECM was initially digested with pepsin, this gap in returned sequences is likely due to the known cleavage sites for pepsin at pH1.3 (pepsin cleavage at 92, 93 and 134 according to PeptideCutter, Swiss Institute for Bioinformatics). These results implicated four peptides of interest, ranging from amino acid residues 55 to 234 of fibrillin-1 (**Table 1**). The first of these peptides, corresponding to residues 55-86 (the largest of the peaks) was synthesized and further validated. Hereafter we refer to this peptide as F1R1

**Table 1:**
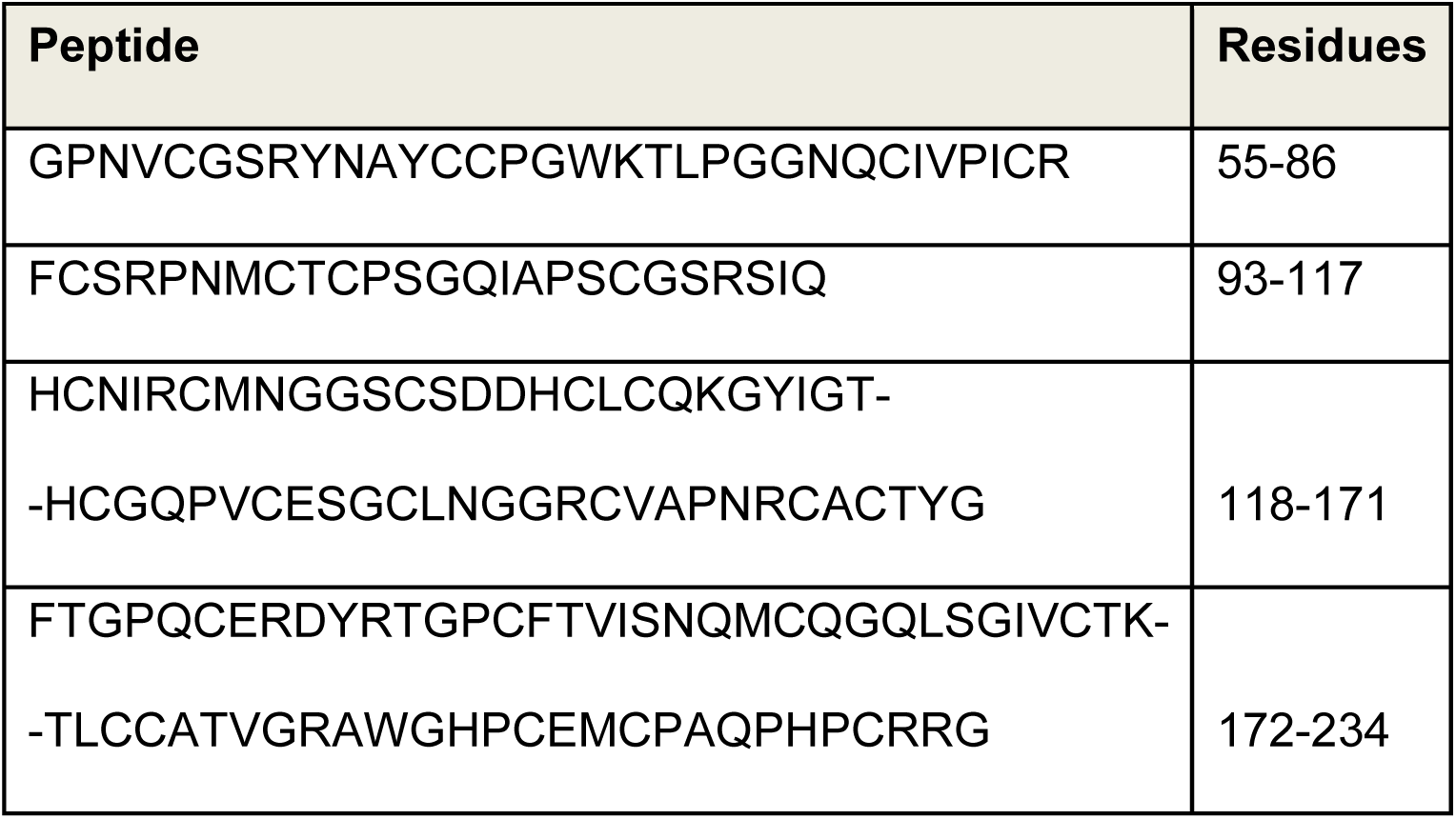
Analysis of the LC-MS/MS peptide sequences in proliferative regions led to the identification of 4 sequences of interest. The first of these sequences we labelled F1R1. Note that the second sequence corresponds to the gap between peaks in **Figure 3B.**

### The F1R1 Tertiary Structure Induces CM Proliferation

To validate the F1R1 peptide’s ability to induce CM proliferation, we conducted a series of in vitro experiments utilizing synthesized F1R1 peptide (GenScript). In the first, F1R1 was synthesized and adsorbed to tissue culture plastic (TCP) at increasing densities between 0 and 12 μg/cm^2^. 6.5μg/cm^2^ was chosen as an approximate mean for adsorption to TCP based on previous experiments indicating 50μg/cm^2^ of total decellularized fetal cECM induced a proliferative response and previously reported fibrillin-1 composition in fetal cECM of ∼13% of the total ECM protein (10). Next, isolated neonatal rat cardiac cells were seeded and cultured on the surfaces (**Figure 4A**). After either 1 or 5 days in culture, the cells were fixed and immunostained for cardiac α-actin, and Ki-67 and imaged with a fluorescent microscope. CM density at 24 hours was not different between any of the groups. At 5 days, CM density was significantly higher for the positive control and all F1R1 groups as compared to their values at 24 hours (**Figure 4B**). In addition, all F1R1 groups displayed a significantly higher CM density at 5 days when compared to the negative control. F1R1 concentrations of 4 μg/cm^2^ and higher also displayed significantly greater CM density than the FBS group after 5 days in culture. In terms of Proliferating CM Density, after 5 days of culture proliferating CM density was also significantly higher on F1R1 adsorbed at 10 μg/cm^2^ than all other groups (**Figure 4C**), while all groups showed a significant increase with time other than the PLL group. The fraction of cells that were CMs was consistent among the groups at 24 hours but significantly increased by day 5 in F1R1 concentrations of 4μg/cm^2^ and higher (**Figure 4D**). Two-way ANOVAs demonstrated a significant interaction between treatment and time for CM Density, Proliferative CM Density and CM Fraction. By comparing cell numbers 24 hours after seeding with those cultured for 5 days, we determined the fold change in CMs to be significantly higher at F1R1 concentrations of 6 and 10 μg/cm^2^ (**Figure 4E**).

**Figure 4:**
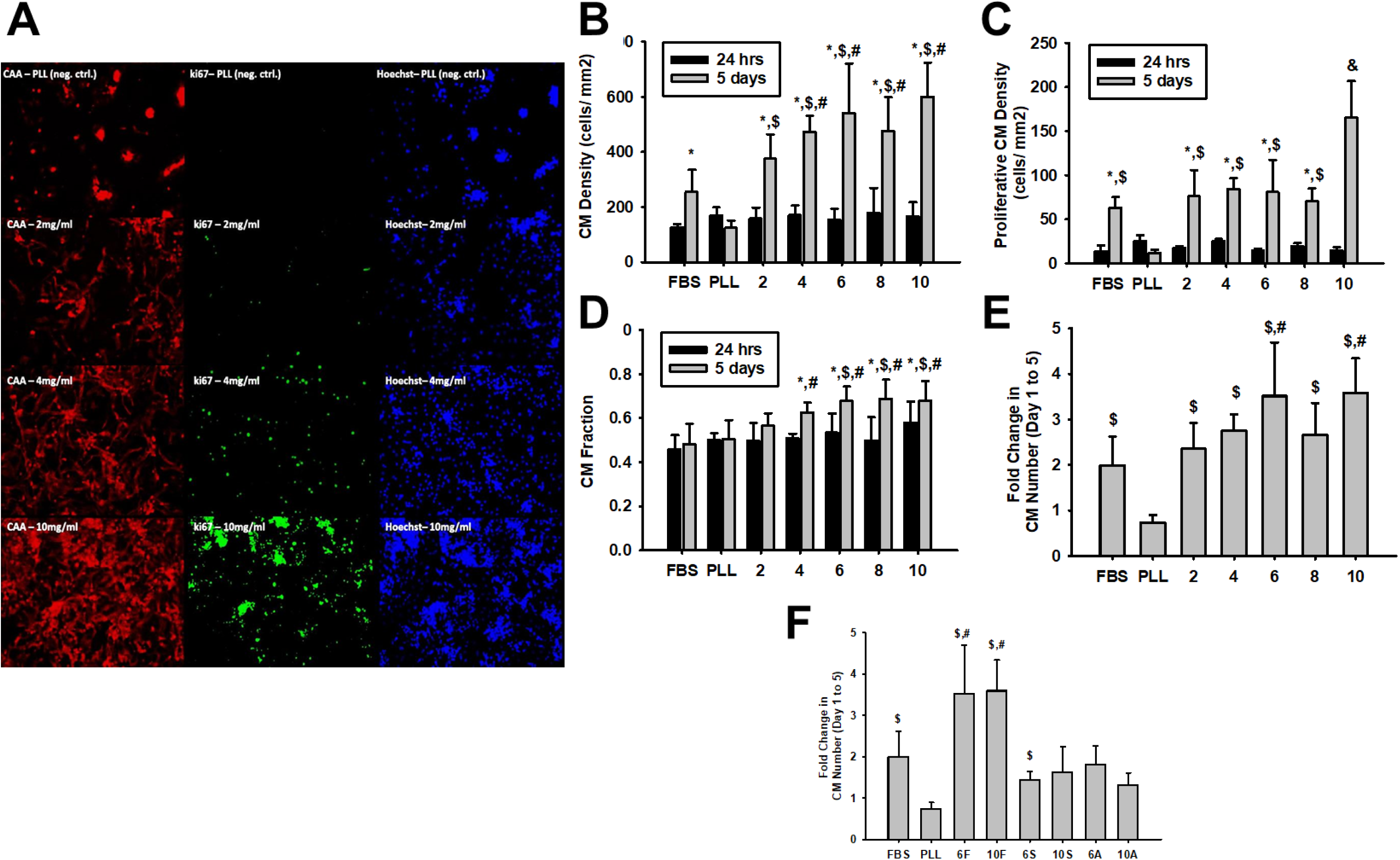
Synthesis of F1R1 and subsequent cell treatment led to cardiomyocyte proliferation at levels similar to whole fetal ECM. (A) Representative images for cardiac a-actin (red, left column), Ki67 (green, middle column) and DAPI (blue, right column) for PLL controls (top row) and 2 μg/cm^2^ (second row), 4 μg/cm^2^ (third row) and 10 μg/cm^2^ (bottom row) F1R1 conditions. Quantified data (n=6 wells per group) for CM Density **(B)**, Proliferative CM Density **(C)** and CM Fraction **(D)** showed increasing CM density, proliferating CM density and CM Fraction with F1R1 treatment compared to PLL and FBS groups in a dose dependent manner. **(E)** Fold change in CM number between 1 day and 5 days in culture showed a significant increase in fold change for all groups compared to PLL and F1R1 doses of 6 and 10 μg/cm^2^ had significantly higher fold change in CM number than FBS controls as well. **(F)** Both scrambled (S) and linearized (A) versions of the F1R1 peptide at the same concentrations showed a loss of the proliferative response, indicating that sequence and tertiary structure are both important in the effect. * denotes p<0.05 w.r.t. 24 hours data for the same group. $ denotes p<0.05 w.r.t. PLL value at the same time point. # denotes p<0.05 w.r.t. FBS value at the same time point. & denotes p<0.05 w.r.t. all other groups at 5-day time point.

In order to further validate this specific sequence was promoting CM proliferation, we repeated the experiment with similar concentrations of a scrambled version of the peptide (add sequence here). While CMs cultured on the F1R1 peptide showed a similar degree of fold change as seen in the previous experiments, those cultured on the scrambled version of the peptide (6S and 10S) were significantly less than the 6 and 10 μg/cm^2^ F1R1 groups and had a similar degree change as the control groups (**Figure 4F**). In addition to the sequence specific effect, we also sought to determine whether the tertiary structure of the F1R1 peptide sequence played a role in promoting CM proliferation. F1R1 contains 5 cysteine residues which should lead to the formation of a tertiary structure via dis-sulfide bond formation. To assess whether this tertiary structure played a role in its effect, the F1R1 peptide was subjected to cysteine alkylation via Iodoacetamide treatment before adsorption to TCP. Alkylation of the cysteine residues prevents folding of the F1R1 peptide leaving a linear version. CMs plated on the linearized alkylated F1R1 (6A and 10A) demonstrated significantly reduced CM proliferation between 1 and 5 days as compared to the comparable concentrations of non-linearized F1R1 (**Figure 4F**).

### Confirmation of a proliferative response in Human iPSC- Derived CMs

In order to determine whether the proliferative effect of the F1R1 peptide was universal and not a response specific to CMs derived from neonatal rats, we also tested its efficacy in human induced pluripotent stem cells derived CMs (iPSC-CMs). Spontaneously beating iPSC-CMs were generated by small molecule differentiation of BJ-RiPS-1.1 induced pluripotent stem cells into cardiomyocytes (25). At day 12 post-differentiation iPSC-CMs were dissociated, enriched with a Percoll gradient, and replated on surfaces coated with either gelatin, gelatin with F1R1 peptide, or gelatin with F1R1-Scramble peptide. EdU (5-ethynyl-2 ’-deoxyuridine) was added to wells at either day 13 or day 19 post-differentiation and incubated for 2-days. At days 15 and 21, the cells were fixed and immunostained for sarcomeric α-Actinin. Fluorescent images were collected and CellProfiler was used to quantify cells with EdU+ nuclei and co-expressing α-Actinin. The iPSC-CMs cultured on the gelatin mixed with F1F1 had a significantly higher fraction of EdU positive CMs as compared to both scrambled F1R1 mixed with gelatin and gelatin alone at both time points (**Figure 5**).

**Figure 5:**
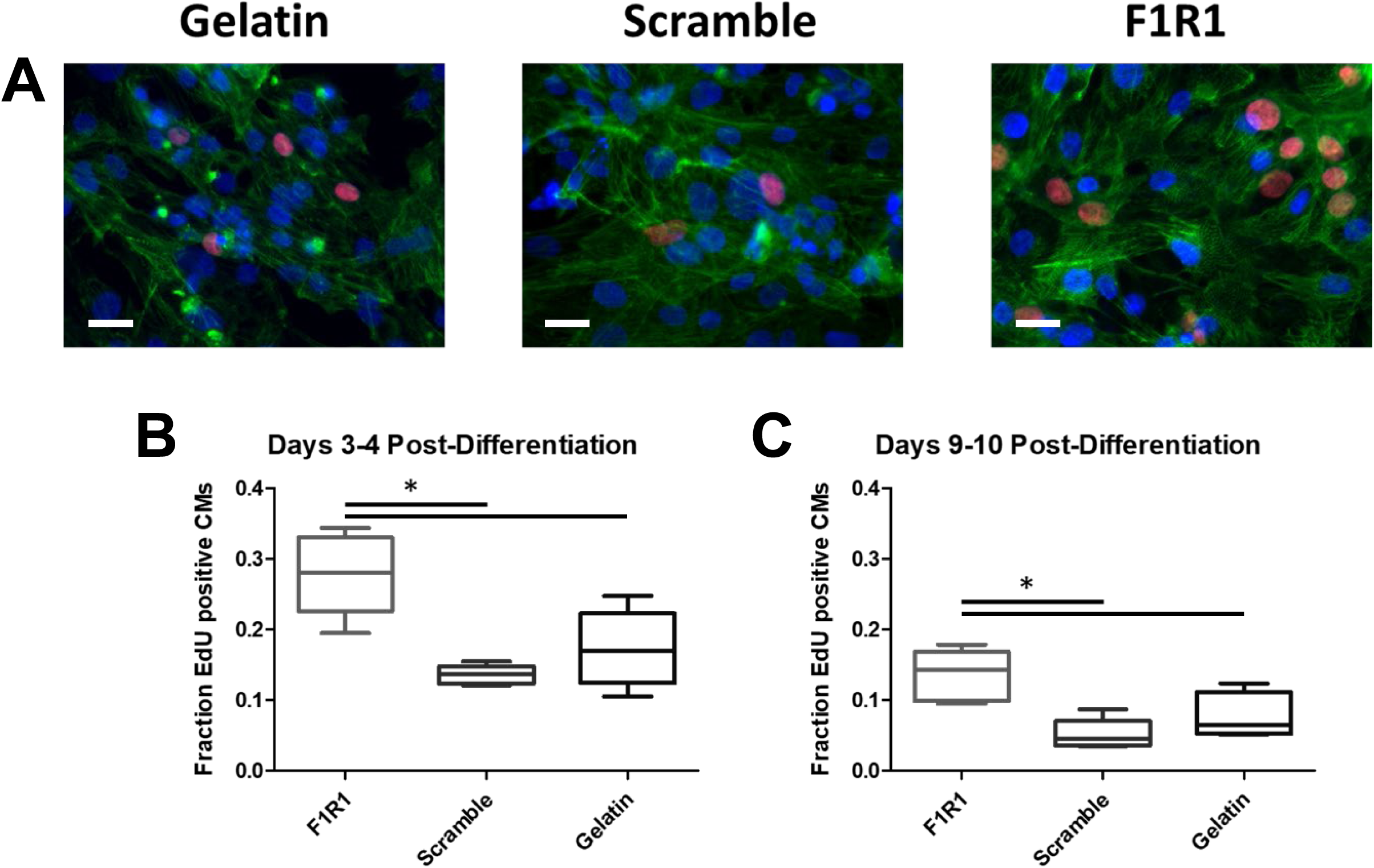
F1R1 stimulated proliferation of human iPSC-derived cardiomyocytes. **(A)** Representative images of human iPSC-derived CMs labeled with EdU (red) for proliferating nuclei, cardiac α-actinin (green) and DAPI (blue) for Gelatin controls (left) as well as 10 μg/cm^2^ Scrambled F1R1 and F1R1 added to gelatin. Quantification of the fraction of EdU positive CMs at both 3-4 days post differentiation **(B)** and 9-10 days post differentiation **(C)** demonstrated a significant increase in proliferating CMs in the F1R1 group as compared to both Scrambled F1R1 and Gelatin controls. * denotes p<0.05.

### Potential mechanism of action from computational analysis of F1R1 sequence

Noting that the tertiary structure and sequence were important to its effects, we sought to use computational methods to further investigate the structure of the F1R1 peptide. In comparing the sequence homology with known proteins, beyond the expected match with fibrillin-1, we also saw a high degree of homology with fibrillin- 2 and fibrillin -3 (**Figure 6A**). Interestingly, we also saw high homology with Latent TGF-β binding protein 2 (LTBP2), particularly with the placement of the 5 cysteine residues- and some homology with LTBP1. Given the homology with LTBP1 and 2, we also used PepSite Finder to assess where the F1R1 peptide would bind with TGF-β1 (**Figure 6B**). These simulations indicated that most likely binding regions for the F1R1 peptide on TGF-β1 were in the Finger 2 region of TGF-β1 which is the critical region for binding between TGF-β1 and the TGF-β receptor 2 (26).

**Figure 6:**
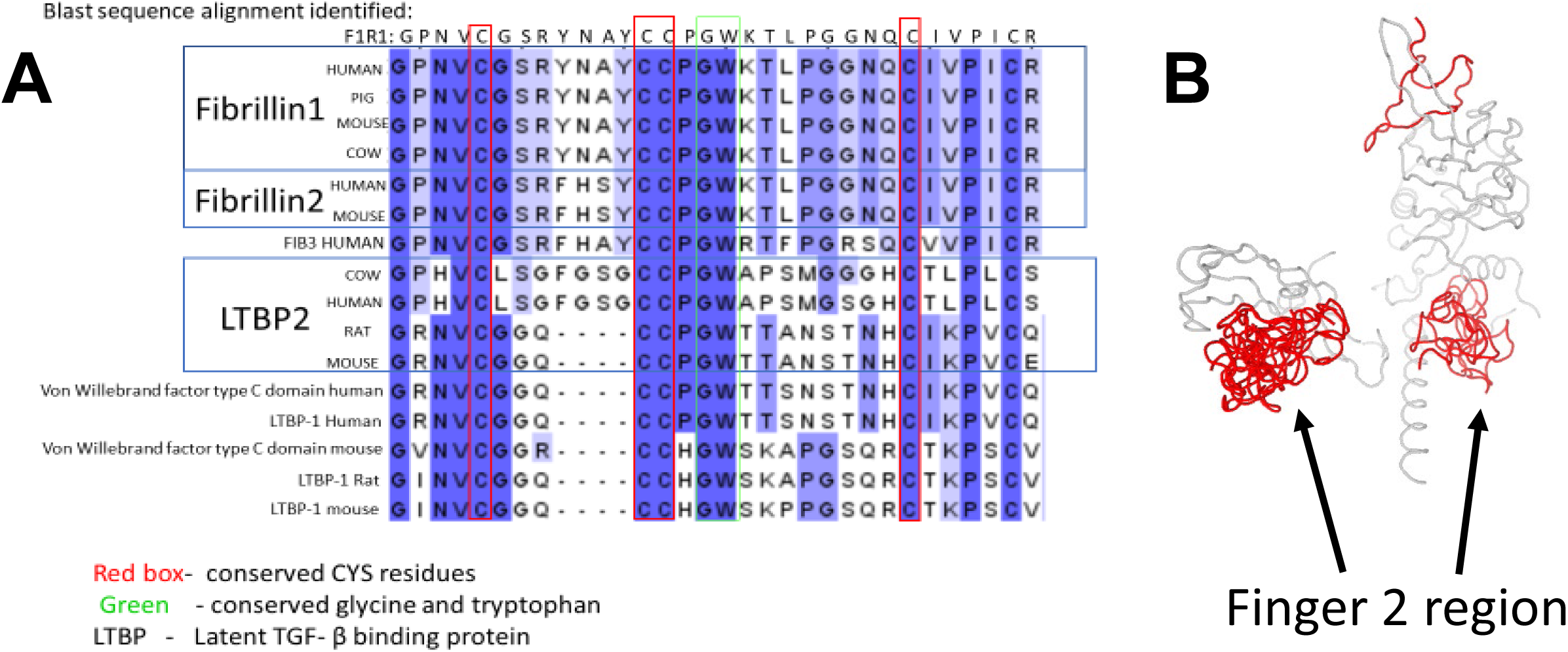
Sequence analysis showed homology with Latent TGF-β binding proteins. **(A)** Sequence analysis carried out in PBLAST demonstrated high sequence homology between F1R1 and fibrillins 2 and 3. In addition, there was relatively well conserved placement of the cysteine residues between F1R1 and latent TGF-β binding peptides 1 and 2, indicating potential structural homology as well. **(B)** PepSite Finder analysis of likely binding sites for F1R1 on TGF-β indicated high likelihood of binding with the Finger 2 region of TGF-β which is critical in binding of TGF-β to TGF-β receptor 2 (F1R1 in red, top 20 results from the simulation shown).

### F1R1 treatment led to reduction in cardiac fibroblast activation

The potential impact of F1R1 on TGF-β signaling was further validated by assessing the impact of F1R1 treatment on cardiac fibroblast activation via TGF-β treatment. The results indicated that while F1R1 did not alter activation levels in the absence of TGF-β1, F1R1 treatment resulted in a significantly reduced percentage of α-Smooth Muscle Actin (α-SMA) positive myofibroblasts in the presence of TGF-β1 as compared to the a commercially available TGF-β1 inhibiting antibody (eBioSciences) and the no treatment control (**Figure 7**). F1R1 treatment was also the only group to show no significant increase in α-SMA+ CFs with the addition of TGF-β1 treatment.

**Figure 7:**
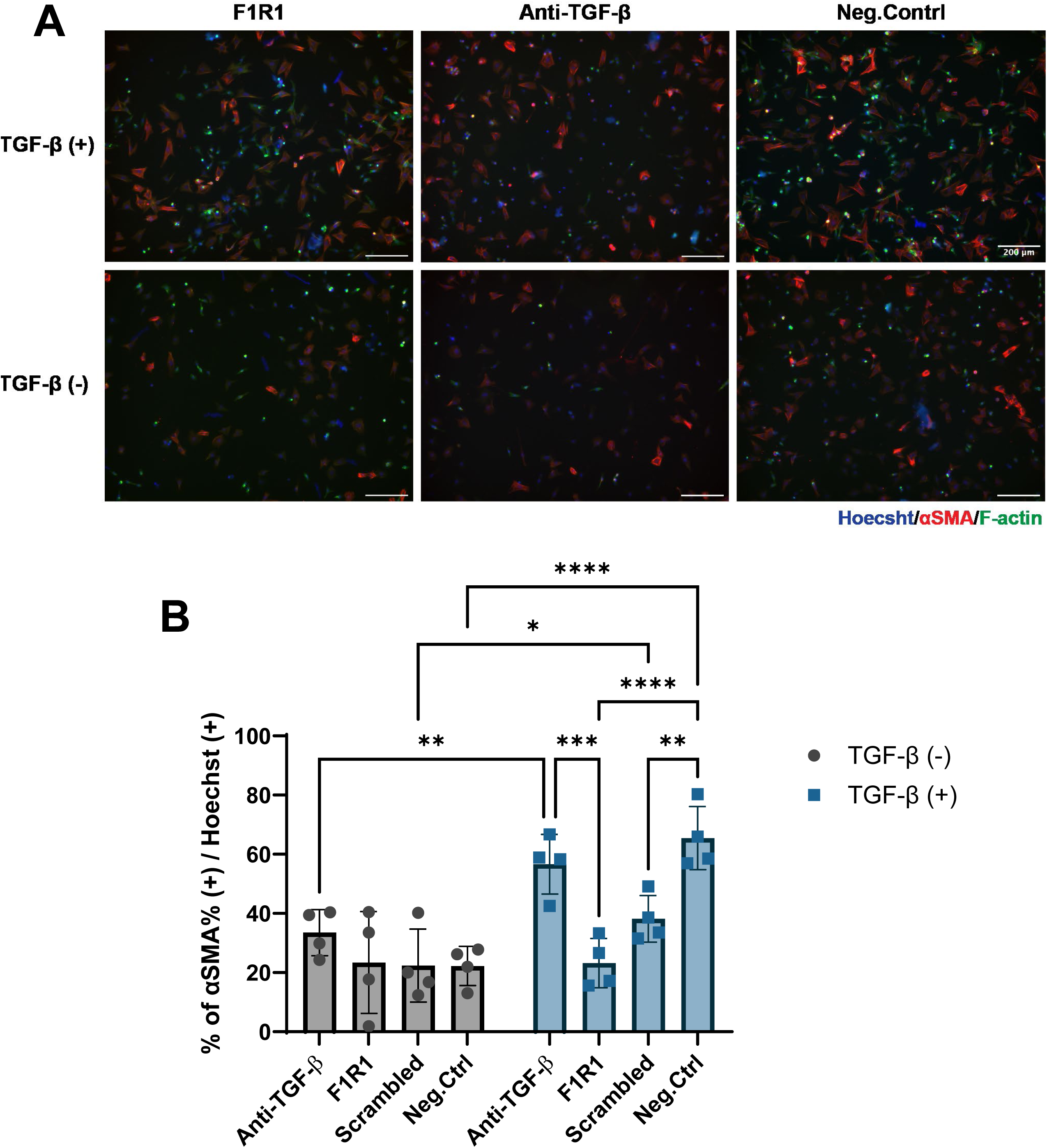
F1R1 treatment significantly reduced cardiac fibroblast activation in response to TGF-β stimulation. **(A)** Representative images of Hoescht (nuclei, blue), α -smooth muscle actin (αSMA, red) and filamentous actin (F-actin, green) labeling of cardiac fibroblast cultures treated with F1R1 (left column), an anti-TGF-β antibody (middle column) or nothing (right column) both with (top row) and without (bottom row) TGF-β stimulation. **(B)** Quantification of images (n=6 wells per group) demonstrated that while there was no difference in activated fibroblasts (% α-SMA+) in the absence of TGF-β stimulation (left), F1R1 treatment led to a significantly lower level of α-SMA+ fibroblasts with TGF-β stimulation and was the only group to not show a significant increase in α-SMA+ fibroblasts with TGF-β treatment. * denotes p<0.05, ** denotes p<0.01, *** denotes p=0.001, **** denotes p<0.001.

## DISCUSSION

Our findings identify a fibrillin-1–derived matrikine, F1R1, as a potent inducer of cardiomyocyte proliferation and highlight a novel strategy for discovering bioactive ECM fragments. By integrating SDS-PAGE fractionation, blot-based functional screening, and LC-MS/MS analysis, we mapped proliferative “hotspots” within solubilized cardiac ECM (**Figure 2**) and pinpointed fibrillin-1 sequences enriched in these regions (**Figure 3**). The ability of synthetic F1R1 to recapitulate the proliferative effect of whole ECM (**Figure 4**) underscores the therapeutic potential of matrikines as defined, sequence-specific agents for cardiac regeneration. Moreover, demonstration that the likely pathway being modulated was related to TGF-b signaling (**Figure 6**) led to the demonstration of reduced cardiac fibroblast activation (**Figure 7**) with F1R1 treatment, highlighting the potential for this method to generate new broad spectrum peptide therapeutics.

Fibrillin-1 is an important structural ECM protein that provides mechanical support and a template for subsequent ECM deposition in many tissues. Fibrillin-1 also plays a role in biochemical cell signaling throughout the body, as Fibrillin-1 microfibrils sequester latent TGF-β (27) and interact with pro-domains of bone morphogenetic proteins (BMPs) (28), where the cysteine-rich nature of fibrillin-1 has been shown to play a large role in the direct sequestration of BMP (29). In addition, the presence of an RGD motif in the fibrillin-1 sequence has implicated its complex involvement in mediating cell binding via integrins (30). In addition, the formation of myofibrils via fibrillins has been cited as having a crucial role in providing strength and regenerative capacities to other organs (31). Finally, mutations in fibrillin-1 result in diseases such as Marfan’s syndrome where improper fibrillin-1 folding results in a lack of TGF-β sequestration, resulting in abhorrent connective tissue growth (32). All of these studies highlight the potential importance of Fibrillin-1 in tissue healing, repair and regeneration.

The proliferative response elicited by our identified F1R1 peptide was robust, with fold-change increases comparable to or exceeding those reported for genetic or pharmacologic interventions targeting cardiomyocyte cell cycle regulators. Proliferation of CMs in the native myocardium has been shown to range from 1.3-4% of the total CM population (33). Studies have reported on a variety of methods for increasing proliferation. Engel et al showed that compared to WT mice, inactivation of the MAP kinase p38α in neonatal CMs led to an increase in the proliferation marker H3 phosphorylation from 0.13% to 0.25% in vivo, a fold change increase of 1.9 (34). Another study by Laflamme et al showed that 14.4% of human CMs from a differentiated cardiac-enriched hESC progeny expressed the Ki-67 marker after 4 weeks in vivo in a rat model of cardiac injury (35). These results are comparable to our findings, where the percent of proliferative CMs after 5 days in controls was 9.5 ± 1.1%, but increased to as much as 27.6 ± 2.1%, with a fold change increase of 2.9 when cultured on F1R1 (10 µg/cm^2^) (**Figure 4E**). In fact, this overall increase in CM proliferation on the F1R1 peptide was similarly apparent, being greater than or equal to 14.8 ± 1.7% in all conditions. In addition, all F1R1 adsorption densities greater than 2 µg/cm^2^ elicited significantly higher overall CM percentages in each well compared to both positive and negative controls (p < 0.05, **Figure 4D**). Importantly, this effect was sequence- and structure-dependent: scrambling the peptide or disrupting its disulfide bonds abolished activity, suggesting that tertiary conformation is critical for biofunction. This structural requirement aligns with fibrillin-1’s known role in microfibril assembly and growth factor sequestration, raising intriguing mechanistic possibilities. Finally, the translational relevance of F1R1 was supported by its activity in human iPSC-derived cardiomyocytes, indicating conservation across species and developmental contexts.

In comparing this with our previous published work using pepsin-solubilized cECM, it is important to note that fibrillin-1 was one of the more abundant proteins present in the fetal cECM that resulted in the highest degree of CM proliferation (10). We had hypothesized in that work that the peptides responsible could either be 1) from ECM proteins that are present in early development but not present in adult heart or 2) present in both the young and adult heart but inhibited or masked by other proteins in the adult heart. In this work we have demonstrated that the adult cECM contains peptides that can promote CM proliferation to the same extent as what is present in fetal cECM (**Figure 4**), supporting the latter of the two hypotheses. Fibrillin-1 has been previously shown to improve the biocompatibility of polytetrafluoroethylene (PTFE) biomaterials (36), promote adhesion, migration and proliferation in human endothelial cells (37), and promote microvascular morphogenesis and wound healing in vitro (38). However, all of this work utilized regions of the protein close to the C-terminus of the fibrillin-1 protein. As such, our identification of the peptides sequence present at the N-terminus of fibrillin-1 present a novel group of peptides that promote cardiomyocyte proliferation and lead to reductions in TGF-β induced cardiac fibroblast activation. This is especially interesting since much of the published work surrounding this region in fibrillin-1 has focused more on growth factor sequestration and its utility in dimerization and fiber formation of the molecule.

Computational modeling revealed homology between F1R1 and latent TGF-β binding protein domains, and in vitro assays demonstrated attenuation of TGF-β–induced fibroblast activation, implicating F1R1 in modulation of profibrotic signaling. LTBP2 also interacts with FGF2 (39), which has been shown to modulate cardiomyocyte proliferation in vitro (40) while also leading to increases in cardiac fibroblast proliferation and activation and subsequent collagen deposition following myocardial injury (41). Together, these data suggest that F1R1 may act through dual mechanisms—direct stimulation of cardiomyocyte proliferation and indirect suppression of adverse remodeling.

Future studies should explore additional fibrillin-1 fragments identified in our proteomic screen, as these regions overlap with EGF-like domains implicated in growth factor interactions. Elucidating the signaling pathways engaged by F1R1—whether via integrins, TGF-β modulation, or other mechanisms such as FGF2 signal modulation—will be critical for rational design of peptide-based therapeutics. Given the structural complexity of ECM signaling, combinatorial strategies incorporating multiple matrikines or pairing peptides with biomaterial scaffolds may offer synergistic benefits for regenerative medicine. Future improvement to this method could include the use of two-dimensional blotting to further fractionate the proteins, which should give a higher fidelity to specific peptides in the fraction. Lastly, beyond cardiac repair, our platform offers a generalizable approach for identifying matrikines that regulate diverse cellular phenotypes, with implications for wound healing, fibrosis, and cancer biology. Future studies could use this same methodology to identify matrikines that are relevant to these biological contexts.

In summary, this work establishes matrikines as a promising class of bioactive molecules and introduces a discovery pipeline that bridges ECM biology and therapeutic innovation. Our work identified a potential therapeutic peptide from whole solubilized ECM that could aid in cardiac repair and regeneration. The fact that we have a peptide that demonstrates similar efficacy levels a whole solubilized ECM in the context of cardiomyocyte proliferation offers even greater evidence for the power of this techniques, as the synthesis of a subset of peptides will be more scalable, consistent and tunable then whole ECM derived products. F1R1 exemplifies how cryptic ECM cues can be harnessed to unlock regenerative potential in the heart and beyond.

## MATERIALS AND METHODS

### Adult and Fetal Rat Heart Harvest, Decellularization, Digestion, and Solubilization

All animal procedures were performed in accordance with the Institutional Animal Care and Use Committee at Tufts University and the NIH Guide for the Care and Use of Laboratory Animals. For adult cECM procurement, approximately 3-month-old pregnant Sprague Dawley rats were anesthetized with a mixture of 100mg/kg ketamine and 10mg/ml xylazine and then euthanized by harvest of the heart. For fetal cECM isolation, fetal pups (embryonic day 18-19) were isolated from the uterus, euthanized by decapitation, and their hearts harvested. Freshly isolated hearts were decellularized according to previously published protocols (10, 18, 19). Adult hearts were decellularized via perfusion decellularization with 1% (wt/vol) sodium dodecyl sulfate (SDS) in deionized water for approximately 48 hours. Fetal hearts were immersed and gently agitated in 0.05% SDS. Next, the cECM was washed for two hours in Triton X-100 (Amresco, Solon, OH) at the same concentration as the respective SDS solution. The cECM was washed three times for ten minutes each with large volumes of deionized water to remove residual detergent. Then the cECM was washed for three days in phosphate-buffered saline (PBS), which was exchanged every 12 hours with the aid of a peristaltic pump. Finally, the ventricular cECM was isolated and the atrial tissues and cardiac skeleton were discarded. The cECM was frozen at −20⁰C, lyophilized, and then mechanically disrupted using microscissors. The cECM was digested and solubilized at room temperature and with constant stirring using 10% w/v pepsin in a 0.1M HCl solution. Adult cECM was digested for approximately 24 hours. Fetal cECM was digested for approximately 1 hour. Following complete solubilization, 1N NaOH was titrated in until the pH was basic (8.5) to quench pepsin activity and then neutralized with 1M HCl to a pH of 7.4.

### cECM SDS-PAGE Fractionation and Transfer to PVDF Membranes

Solubilized cECM peptides were fractionated by molecular weight using standard sodium dodecyl sulphate–polyacrylamide gel electrophoresis (SDS-PAGE) methods for cellular protein separation. First, the solubilized cECM solution was sonicated on ice with 20 second pulses at 30% amplitude (Branson Digital Sonifer) followed by denaturization which was achieved by heating the samples to 70C for 10 min. The samples were then treated with DTT to reduce the proteins for subsequent PAGE. Prepared samples were intentionally loaded onto a 10% polyacrylamide gel in a precise order to create mirror images on the left and right halves gel. (e.g. Ladder, sample 1,… sample 4, sample 4, … sample 1, ladder). The gel was run at 100V at room temperature until the dye front reached the bottom of the gel (∼50-60 minutes). After electrophoresis, half of the gel was transferred overnight at 75 volts at 4°C to a polyvinyl difluoride (PVDF) membrane (Biorad). The second half of the gel was kept for extraction of peptides and subsequent LC-MS/MS compositional analysis. The transferred membrane was sterilized via three rinses of 1% penicillin-streptomycin (Invitrogen) in PBS for 15 minutes each and then blocked in 1% bovine serum albumin (BSA). The membrane was then transferred to a sterile tissue culture dish with the side containing the transferred fractionated cECM facing upwards.

### Primary Neonatal Rat Cardiac Cell Isolation and Culture

P2-P3 neonatal Sprague Dawley rat pups were euthanized by conscious decapitation. Hearts were harvested and tissue minced. Minced tissue was digested 7 times, for 7 mins each, with collagenase type 2 in sterile PBS supplemented with 20mM glucose. Cells were strained through a 70µM filtered and seeded onto various culture surfaces as described for each specific experiment. For all experiments, cells were cultured in media composed of a 50/50 mixture of DMEM and Ham’s F12 Nutrient Mix (Invitrogen), 10% horse serum, 2% fetal bovine serum, 0.5% Insulin-Transferrin-Selenium-X (Invitrogen), and 1% penicillin-streptomycin (Invitrogen) (10). For serum free medias, 0.2% BSA (wt/vol) (Sigma) was used instead of horse and bovine serums.

### Immunocytochemistry and Image Acquisition

For all immunocytochemistry analysis cells were fixed with 4% paraformaldehyde at 4°C for 20 minutes and permeabilized with 0.1% Triton X-100 for 20°C for 10 minutes. Cells were blocked with 5% donkey serum and 1% BSA (Sigma) in PBS. Next, cells were sequentially incubated with anti-α-actin and anti-Ki67 antibodies and then co-incubated with Alexa Fluor 555 anti-mouse and Alexa Fluor 488 anti-rabbit secondary antibodies. Details of antibodies used are in **Table 2**. All incubation steps were conducted at 20⁰C for 1 hour. Nuclei were stained by incubation with Hoechst 33258 (Invitrogen). Immunostained cells were imaged using an Olympus IX-81 inverted fluorescence microscope (Olympus America, Center Valley, PA). Image acquisition was automated using a Prior stage and the multidimensional acquisition settings in Metamorph (version 7.7.4.0, Molecular Devices, Inc., Sunnyvale, CA.).

**Table 2:** Antibodies used to immunostaining proliferating cardiomyocytes cultured on PVDF membranes blotted with fractioned cECM.

| Target | Host | Conjugation | Manufacturer | Product # |
| --- | --- | --- | --- | --- |
| anti-α-actin | mouse | - | Santa Cruz | Sc-58670 |
| anti-Ki67 | rabbit | - | Santa Cruz | sc-23900 |
| anti-rabbit | donkey | Alexa Fluor® 488 | Jackson<br>ImmunoResearch | 711-545-152 |
| anti-mouse | donkey | Alexa Fluor® 555 | Invitrogen | A32773 |

### Detection of Proliferative CM Regions within the Blot Culture

Freshly isolated neonatal rat cardiac cells were seeded at 100,000cells/cm^2^ directly onto the surface of the PVDF membrane with the fractionated cECM. Cells were cultured in serum free media and the media changed every other day. On day 3 or 5, the cells were fixed, permeabilized, blocked, stained for nuclei, cardiac α-actin, and Ki-67, and fluorescent images acquired as described in the ‘*Immunocytochemistry and Image Acquisition’* section. After acquisition, the entire PVDF membrane was reconstructed by aligning sequential fluorescent images. The reconstructed image was analyzed with a custom CellProfiler (r11710, The Broad Institute, Cambridge, MA) pipeline to determine the total numbers of cell nuclei, cardiac α-actin+ cells, and Ki-67+ cells as a function of molecular weight.

### Proteomic Analysis for Identification of cECM Peptides that Promotes CM Proliferation

Molecular weight regions that corresponded to increased CM proliferation on the blot culture were excised from the other saved half of the gel with a razor blade. Regions with little to no CM proliferation were also excised and used as controls. Gel fragments were digested at a concentration of 5mg/ml in a solution containing 5M urea, 2M thiourea, 50mM DTT and 0.1% SDS in PBS (42) with constant stirring at 4⁰C for 48 hours. Afterwards, the samples were sonicated on ice with 20 sec pulses at 30% amplitude (Branson Digital Sonifer) and the cECM peptides were precipitated with acetone. Samples were then prepped for for liquid chromatography tandem mass spectrometry (LC-MS/MS) analysis as previously described (ref) and spectra were mapped to rat proteins using a standard library in ProteinPilot (SCIEX, Marlborough, MA). Analysis of the relative abundance of each protein present in the proliferative or control exicision region was carried out by normalizing the number of spectra for a given protein by the total number of ECM protein spectra identified in the analysis. In order to determine which peptides were most prevalent within gel fragments corresponding to increased levels of CM proliferation, a custom program written in the C# language (Microsoft) was used to load all the obtained peptide sequences from LC-MS/MS and return an amino acid residue histogram, with peptide sequences ordered by both greatest length and frequency of occurrence. By assessing sequences present in proliferative regions that were not in control regions, we were able to identify several potential candidate peptide sequences that promoted CM proliferation.

### Assessment of CM Adhesion and Proliferation on Synthetic F1R1 Peptide Surfaces

Following analysis of the peptide sequences most present in proliferative regions of the blot, we identified a region of the ECM protein fibrillin-1 that appeared to be present in higher-than-expected abundance in proliferative regions of the blot culture. The F1R1 peptide (Sequence: GPNVCGSRYNAYCCPGWKTLPGGNQCIVPICR) and a scrambled version of the F1R1 peptide (Sequence: VPCNLSCQINAYYCWGPKTVRGGCPGCNPGIR) were synthetically manufactured (95% purity, TFA removed) (GenScript, Piscataway, NJ) and allowed to adsorb to the surface of TCP by drying at room temperature overnight in a sterile tissue culture cabinet. F1R1 was adsorbed at 2, 4, 6, 8, and 10µg/cm^2^. Scrambled and linearized (see ‘*F1R1 Peptide Alkylation’ section)* versions of the peptide were adsorbed at concentrations of 6 and 10 µg/cm^2^. Non-absorbed TCP and serum containing media was used as a positive control. For a negative control, TCP was incubated wit 100µl of a 0.01% poly-l-lysine (PLL) solution for 5 minutes at room temperature. Once dry, surfaces were rinsed twice with sterile PBS and neonatal rat cardiac cells, isolated as described in the *‘Primary Neonatal Rat Cardiac Cell Isolation and Culture’* section, were seeded at 10,000 cells/cm^2^ onto the surfaces. Experimental and negative control groups were cultured in serum free media and the media changed every other day. On day 1 or 5, the cells were fixed, permeabilized, blocked, stained for nuclei, cardiac α-actin, and Ki-67, and fluorescent images acquired as described in the ‘*Immunocytochemistry and Image Acquisition’* section. The images were analyzed with a custom CellProfiler (r11710, The Broad Institute, Cambridge, MA) pipeline to determine the total numbers of cell nuclei, cardiac α-actin+ cells, and Ki-67+ cells.

### F1R1 Peptide Alkylation

To assess whether the tertiary structure of the F1R1 peptide was important in the cellular response to it, we sought to linearize the peptide by alkylating the cysteine residues prior to adsorption to TCP. To do this, 500μg of the peptide was resuspended in 500μl of tris/urea buffer (9M urea, 40 mm Tris, pH 8.3) supplemented with 50mm DTT and incubated at 37°C for 30 minutes to reduce disulfide bonds. 75μl of 0.5M iodoacetamide was then added to alkylate cysteine residues, and the solution was left to incubate for 30 minutes in the dark at room temperature. The alkylated peptide was then precipitated with ice cold acetone for 1 hour at −20°C and centrifuged at 13,000rpm for 25 minutes. The pellet was washed once with acetone, and the supernatant was aspirated leaving the pellet to air-dry before being resuspended in PBS. The alkylated peptide was then subjected to a BCA assay to determine the new peptide concentration before adsorbing to TCP as described above.

### Proliferation of Human iPS-Derived Cardiomyocytes on Synthetic F1R1 Peptide Surfaces

Human BJ-RiPS-1.1 cells were obtained from the Harvard Stem Cell Institute and cultured on 1/100 diluted growth factor reduced MatrigelTM (Cat. CB 40230, Thermo Fisher Scientific Inc) in mTeSRTM1 media (Cat. 05870, Stem Cell Technologies Inc). Cultures were maintained between passage 18-30. Cardiac differentiation was performed using an established protocol based on a temporal modulation of Wnt signaling with small molecules (Stemolecule™ CHIR99021, Cat. 04-0004-02; Stemolecule™ Wnt Inhibitor IWP-4, Cat. 04-0036; Stemgent Inc), generating spontaneously beating cardiomyocytes by day 10-12 post-differentiation (25). Cardiomyocytes were maintained in RPMI 1640 medium (Cat. 11875-119, Life Technologies) supplemented with B-27 (Cat. 17504-044, Life Technologies), with media changes every 2 days. At day 12, cardiomyocytes were dissociated, resuspended in RPMI/B27 medium, and enriched using a Percoll gradient (43). Briefly, a discontinuous gradient was established with a 40.5% Percoll layer, a 58.5% Percoll layer, and a third dissociated cells layer. The gradient layers were centrifuged at 1500g for 30 minutes. The densest fraction (fraction IV) containing a higher percentage of cardiomyocytes was collected, washed, and used for experimentation.

24-well plates were coated with either gelatin, gelatin with 10µg/cm^2^ F1R1 peptide, or gelatin with 10µg/cm^2^ F1R1-Scramble peptide and enriched cardiomyocytes were replated at a density of 25,000 cells/well. EdU (5-ethynyl-2 ’-deoxyuridine) was added to wells at either day 13 or day 19 post-differentiation and incubated for 2-days. At days 15 and 21, cardiomyocytes were fixed with 4% paraformaldehyde at 4°C, washed with PBS, permeabilized with 1.5% Triton-X solution for 10 minutes, washed with PBS, and then blocked with 1.5% donkey serum for 30 minutes. Following blocking, the fixed cells were incubated with anti-sarcomeric α-Actinin antibody (1:100; Cat. A8711, Sigma-Aldrich) overnight at 4°C. Anti-mouse 488 secondary antibody (1:400, Invitrogen) was incubated for 1hr. EdU incorporation was detected using a Click-iT® EdU Imaging Kit with an Alexa Fluor® 594 probe (Cat. C10339, Invitrogen), per the manufacturer’s protocol. Cell nuclei were stained with Hoechst dye (Cat. H3570, Thermo Fisher Scientific, Inc.), washed three times with PBS, and imaged in PBS. For analysis, 20x images (n=5 for each group, at both time points) were collected using a Nikon Eclipse TE200 microscope. A CellProfiler image analysis pipeline and CellProfiler Analyst were used to quantify EdU+ nuclei co-expressing α-Actinin, with 81.40 ± 19.14 total cells/field for day 15, and 114.40 ± 30.32 total cells/field for day 21.

### Computational analysis of F1R1 structural homology with other proteins

In order to assess the structure of the F1R1 peptide we first looked at sequence homology using BLAST. The algorithm of this tool is used to identify local similarities between different nucleic or peptide sequences against their corresponding databases. It then calculates statistical significance to identify best matches. For our purpose, we used BLASTP to identify proteins sequences and UNIPROT to identify which protein our peptide sequence was derived from. To assess where on the TGF-β macromolecule the F1R1 peptide would bind, we used PEP-SiteFinder (44) and mapped the top 20 potential binding locations for F1R1 on TGF-β

### Cardiac Fibroblast treatment with F1R1

Primary neonatal cardiac fibroblasts (nCFs) were isolated from Sprague Dawley rat pups (P2) as previously described (24) and seeded onto a soft substrate (5% wt/v GelMA) at 50,000 cells/cm2 to avoid activation. After pre-plating (45), CFs were starved with serum-free media (50/50 mixture of DMEM and Ham’s 12 Nutrient Mix (Invitrogen) with the supplement of 0.2% bovine serum (BSA) (wt/v)(Sigma), 0.5% Insulin-Transferrin-Selenium-X (Invitrogen), and 1% penicillin-streptomycin (Invitrogen) for 24 hrs. The nCFs were then pre-stimulated with each condition for 12 hrs (F1R1 peptide (1.67 mg/mL, 95% purity, GenScript), scrambled F1R1 (1.67 mg/mL, GenScript), TGF-β neutralizing antibody (50 µg/mL, eBioscienceTM), and nothing as a negative control) without the presence of TGF-β. TGF-β protein (10 ng/mL, Sino Biological) was then added to the serum-free media and mixed with each cultural condition to treat the nCFs for 24 hours. Cells were then fixed with 4% PFA and stained with Hoechst (nuclei), ɑ-smooth muscle actin (ɑ-SMA, Invitrogen), and phalloidin (F-actin, Invitrogen) in order to assess myofibroblast activation. The fluorescent images were acquired using a Keyence 710 and analyzed with Fiji, ImageJ, and CellProfiler for the quantified results.

### Statistical Analysis

Statistical analysis was performed using appropriately sized ANOVA with posthoc testing carried out using a Tukey’s T-test. Differences were considered statistically significant for p < 0.05. Sample sizes are reported for each data set.

## AUTHOR CONTRIBUTIONS

KJE was responsible for design and conducting the majority of the experiments with F1R1 and cardiomyocytes and writing portions of the manuscript. ECP was responsible for assisting with the experiments and drafting and editing this manuscript. YL carried out the experiments and data analysis related to the cardiac fibroblast study. JG carried out experiments and data analysis related to the iPSC-CM study. AJ carried out analysis related to the sequence homology and binding locations of F1R1. CW assisted in experimental design and data interpretation of the manuscript. JW assisted in experimental design and data interpretation of the manuscript. HO assisted in experimental design and data interpretation of the manuscript and funded the studies on iPSC-CMs. LB assisted in experimental design and data interpretation and writing of the manuscript as well as funding all experiments other than the iPSC-CM studies.

## ACKNOWLEDGEMENTS

The authors would like to acknowledge the assistance of the BIDMC-Harvard Mass Spectrometry Core Facility and its director John Asara for assistance with running and analyzing the proteomics data. In addition, the authors would like to acknowledge funding from the NIH (R21HL115570, R01HL171396, and R56HL153984 all to LB), the Department of Defense (W81XWH-16-1-0304 to LB), the National Science Foundation (NSF1603524 to LB) and the American Heart Association (20TPA35500082 to LB).

## ABBREVIATIONS

BSA: bovine serum albumin
BCA: bicinchoninic acid
BMP: bone morphogenic protein
CHD: congenital heart defect
DTT: dithiothreitol
ECM: extracellular matrix
cECM: cardiac extracellular
matrix CM: cardiomyocyte
iPSC-CMs: induced pluripotent derived cardiomyocytes
LC-MS/MS: liquid chromatography - tandem mass spectrometry
RNA: ribonucleic acid
NIH: National Institutes of Health
PBS: phosphate buffered saline
PLL: poly-l-lysine
PTFE: polytetrafluoroethylene
PVDF: polyvinylidene difluoride
SDS: sodium dodecyl sulfate
SDS-PAGE: sodium dodecyl sulphate–polyacrylamide gel electrophoresis
TCP: tissue culture plastic
TGF-β: transforming growth factor Beta

## REFERENCES

1. Hynes, R.O., The extracellular matrix: not just pretty fibrils. Science, 2009. 326(5957): p. 1216–9, PMCID: PMC3536535, PMID: 19965464

2. Engler, A.J., S. Sen, H.L. Sweeney, and D.E. Discher, Matrix elasticity directs stem cell lineage specification. Cell, 2006. 126(4): p. 677–89, PMCID: PMID: 16923388

3. Ricard-Blum, S. and R. Salza, Matricryptins and matrikines: biologically active fragments of the extracellular matrix. Exp Dermatol, 2014. 23(7): p. 457–63, PMCID: PMID: 24815015

4. Bellon, G., L. Martiny, and A. Robinet, Matrix metalloproteinases and matrikines in angiogenesis. Crit Rev Oncol Hematol, 2004. 49(3): p. 203–20, PMCID: PMID: 15036261

5. Akthar, S., D.F. Patel, R.C. Beale, T. Peiro, X. Xu, A. Gaggar, P.L. Jackson, J.E. Blalock, C.M. Lloyd, and R.J. Snelgrove, Matrikines are key regulators in modulating the amplitude of lung inflammation in acute pulmonary infection. Nat Commun, 2015. 6: p. 8423, PMCID: PMC4595997, PMID: 26400771

6. Jariwala, N., M. Ozols, M. Bell, E. Bradley, A. Gilmore, L. Debelle, and M.J. Sherratt, Matrikines as mediators of tissue remodelling. Adv Drug Deliv Rev, 2022. 185: p. 114240, PMCID: PMID: 35378216

7. Vivien, C.J., J.E. Hudson, and E.R. Porrello, Evolution, comparative biology and ontogeny of vertebrate heart regeneration. NPJ Regen Med, 2016. 1: p. 16012, PMCID: PMC5744704, PMID: 29302337

8. Zheng, L., Y. Chen, and J.W. Xiong, Rewiring cell identity and metabolism to drive cardiomyocyte proliferation. Cell Regen, 2025. 14(1): p. 40, PMCID: PMC12477098, PMID: 41015973

9. Porrello, E.R., A.I. Mahmoud, E. Simpson, J.A. Hill, J.A. Richardson, E.N. Olson, and H.A. Sadek, Transient regenerative potential of the neonatal mouse heart. Science, 2011. 331(6020): p. 1078–80, PMCID: 3099478, PMID: 21350179

10. Williams, C., K.P. Quinn, I. Georgakoudi, and L.D. Black, 3rd, *Young developmental age cardiac extracellular matrix promotes the expansion of neonatal cardiomyocytes in vitro*. Acta Biomater, 2014. 10(1): p. 194–204, PMCID: PMC3840040, PMID: 24012606

11. Bassat, E., Y.E. Mutlak, A. Genzelinakh, I.Y. Shadrin, K. Baruch Umansky, O. Yifa, D. Kain, D. Rajchman, J. Leach, D. Riabov Bassat, Y. Udi, R. Sarig, I. Sagi, J.F. Martin, N. Bursac, S. Cohen, and E. Tzahor, The extracellular matrix protein agrin promotes heart regeneration in mice. Nature, 2017. 547(7662): p. 179–184, PMCID: PMC5769930, PMID: 28581497

12. Kuhn, B., F. del Monte, R.J. Hajjar, Y.S. Chang, D. Lebeche, S. Arab, and M.T. Keating, Periostin induces proliferation of differentiated cardiomyocytes and promotes cardiac repair. Nat Med, 2007. 13(8): p. 962–9, PMCID: PMID: 17632525

13. Bersell, K., S. Arab, B. Haring, and B. Kuhn, Neuregulin1/ErbB4 signaling induces cardiomyocyte proliferation and repair of heart injury. Cell, 2009. 138(2): p. 257–70, PMCID: PMID: 19632177

14. Frangogiannis, N.G., The extracellular matrix in myocardial injury, repair, and remodeling. J Clin Invest, 2017. 127(5): p. 1600–1612, PMCID: PMC5409799, PMID: 28459429

15. Holle, A.W. and A.J. Engler, More than a feeling: discovering, understanding, and influencing mechanosensing pathways. Curr Opin Biotechnol, 2011. 22(5): p. 648–54, PMCID: PMC3150613, PMID: 21536426

16. Miao, M.Z., J.S. Lee, K.M. Yamada, and R.F. Loeser, Integrin signalling in joint development, homeostasis and osteoarthritis. Nat Rev Rheumatol, 2024. 20(8): p. 492–509, PMCID: PMC11886400, PMID: 39014254

17. Leroux, R., C. Ringenbach, T. Marchand, O. Peschard, P. Mondon, and P. Criton, A new matrikine-derived peptide up-regulates longevity genes for improving extracellular matrix architecture and connections of dermal cell with its matrix. Int J Cosmet Sci, 2020. 42(1): p. 53–59, PMCID: PMID: 31596957

18. Ott, H.C., T.S. Matthiesen, S.K. Goh, L.D. Black, S.M. Kren, T.I. Netoff, and D.A. Taylor, Perfusion-decellularized matrix: using nature’s platform to engineer a bioartificial heart. Nat Med, 2008. 14(2): p. 213–21, PMCID: PMID: 18193059

19. Singelyn, J.M., J.A. DeQuach, S.B. Seif-Naraghi, R.B. Littlefield, P.J. Schup-Magoffin, and K.L. Christman, Naturally derived myocardial matrix as an injectable scaffold for cardiac tissue engineering. Biomaterials, 2009. 30(29): p. 5409–16, PMCID: PMC2728782, PMID: 19608268

20. Silva, A.C., S.C. Rodrigues, J. Caldeira, A.M. Nunes, V. Sampaio-Pinto, T.P. Resende, M.J. Oliveira, M.A. Barbosa, S. Thorsteinsdottir, D.S. Nascimento, and O.P. Pinto-do, Three-dimensional scaffolds of fetal decellularized hearts exhibit enhanced potential to support cardiac cells in comparison to the adult. Biomaterials, 2016. 104: p. 52–64, PMCID: PMID: 27424216

21. Wang, Z., D.W. Long, Y. Huang, W.C.W. Chen, K. Kim, and Y. Wang, Decellularized neonatal cardiac extracellular matrix prevents widespread ventricular remodeling in adult mammals after myocardial infarction. Acta Biomater, 2019. 87: p. 140–151, PMCID: PMC6414086, PMID: 30710713

22. Williams, C., K. Sullivan, and L.D. Black, 3rd, *Partially Digested Adult Cardiac Extracellular Matrix Promotes Cardiomyocyte Proliferation In Vitro*. Adv Healthc Mater, 2015. 4(10): p. 1545–54, PMCID: PMC4504755, PMID: 25988681

23. Erin, E.M., G.R. Jenkins, O.M. Kon, A.S. Zacharasiewicz, G.C. Nicholson, H. Neighbour, R.C. Tennant, A.J. Tan, B.R. Leaker, A. Bush, P.J. Jose, P.J. Barnes, and T.T. Hansel, Optimized dialysis and protease inhibition of sputum dithiothreitol supernatants. Am J Respir Crit Care Med, 2008. 177(2): p. 132–41, PMCID: PMID: 17962642

24. Morgan, K.Y. and L.D. Black, 3rd, *Mimicking isovolumic contraction with combined electromechanical stimulation improves the development of engineered cardiac constructs*. Tissue Eng Part A, 2014. 20(11-12): p. 1654–67, PMCID: 4029049, PMID: 24410342

25. Lian, X., J. Zhang, S.M. Azarin, K. Zhu, L.B. Hazeltine, X. Bao, C. Hsiao, T.J. Kamp, and S.P. Palecek, Directed cariomyocyte differentiation from human pluripotent stem cells by modulating Wnt/beta-catenin signaling under fully defined conditions. Nature Protocols, 2013. 8(1): p. 162–175, PMCID: PMID:

26. Qian, S.W., J.K. Burmester, M.L. Tsang, J.A. Weatherbee, A.P. Hinck, D.J. Ohlsen, M.B. Sporn, and A.B. Roberts, Binding affinity of transforming growth factor-beta for its type II receptor is determined by the C-terminal region of the molecule. J Biol Chem, 1996. 271(48): p. 30656–62, PMCID: PMID: 8940041

27. Ono, R.N., G. Sengle, N.L. Charbonneau, V. Carlberg, H.P. Bachinger, T. Sasaki, S. Lee-Arteaga, L. Zilberberg, D.B. Rifkin, F. Ramirez, M.L. Chu, and L.Y. Sakai, Latent transforming growth factor beta-binding proteins and fibulins compete for fibrillin-1 and exhibit exquisite specificities in binding sites. J Biol Chem, 2009. 284(25): p. 16872–16881, PMCID: PMC2719323, PMID: 19349279

28. Sengle, G., N.L. Charbonneau, R.N. Ono, T. Sasaki, J. Alvarez, D.R. Keene, H.P. Bachinger, and L.Y. Sakai, Targeting of bone morphogenetic protein growth factor complexes to fibrillin. J Biol Chem, 2008. 283(20): p. 13874–88, PMCID: PMC2376219, PMID: 18339631

29. Ramirez, F. and L.Y. Sakai, Biogenesis and function of fibrillin assemblies. Cell Tissue Res, 2010. 339(1): p. 71–82, PMCID: PMC2819175, PMID: 19513754

30. Pfaff, M., D.P. Reinhardt, L.Y. Sakai, and R. Timpl, Cell adhesion and integrin binding to recombinant human fibrillin-1. FEBS Lett, 1996. 384(3): p. 247–50, PMCID: PMID: 8617364

31. Olivieri, J., S. Smaldone, and F. Ramirez, Fibrillin assemblies: extracellular determinants of tissue formation and fibrosis. Fibrogenesis Tissue Repair, 2010. 3: p. 24, PMCID: PMC3012016, PMID: 21126338

32. Judge, D.P. and H.C. Dietz, Marfan’s syndrome. Lancet, 2005. 366(9501): p. 1965–76, PMCID: PMC1513064, PMID: 16325700

33. Malliaras, K. and J. Terrovitis, Cardiomyocyte proliferation vs progenitor cells in myocardial regeneration: The debate continues. Glob Cardiol Sci Pract, 2013. 2013(3): p. 303–15, PMCID: PMC3963760, PMID: 24689031

34. Engel, F.B., M. Schebesta, M.T. Duong, G. Lu, S. Ren, J.B. Madwed, H. Jiang, Y. Wang, and M.T. Keating, p38 MAP kinase inhibition enables proliferation of adult mammalian cardiomyocytes. Genes Dev, 2005. 19(10): p. 1175–87, PMCID: PMC1132004, PMID: 15870258

35. Laflamme, M.A., J. Gold, C. Xu, M. Hassanipour, E. Rosler, S. Police, V. Muskheli, and C.E. Murry, Formation of human myocardium in the rat heart from human embryonic stem cells. Am J Pathol, 2005. 167(3): p. 663–71, PMCID: PMC1698736, PMID: 16127147

36. Hajian, H., S.G. Wise, D.V. Bax, A. Kondyurin, A. Waterhouse, L.L. Dunn, C.M. Kielty, Y. Yu, A.S. Weiss, M.M. Bilek, P.G. Bannon, and M.K. Ng, Immobilisation of a fibrillin-1 fragment enhances the biocompatibility of PTFE. Colloids Surf B Biointerfaces, 2014. 116: p. 544–52, PMCID: PMID: 24572497

37. Mariko, B., Z. Ghandour, S. Raveaud, M. Quentin, Y. Usson, J. Verdetti, P. Huber, C. Kielty, and G. Faury, Microfibrils and fibrillin-1 induce integrin-mediated signaling, proliferation and migration in human endothelial cells. Am J Physiol Cell Physiol, 2010. 299(5): p. C977–87, PMCID: PMID: 20686071

38. Demidova-Rice, T.N., A. Geevarghese, and I.M. Herman, Bioactive peptides derived from vascular endothelial cell extracellular matrices promote microvascular morphogenesis and wound healing in vitro. Wound Repair Regen, 2011. 19(1): p. 59–70, PMCID: PMC3059781, PMID: 21134032

39. Menz, C., M.K. Parsi, J.R. Adams, M.A. Sideek, Z. Kopecki, A.J. Cowin, and M.A. Gibson, LTBP-2 Has a Single High-Affinity Binding Site for FGF-2 and Blocks FGF-2-Induced Cell Proliferation. PLoS One, 2015. 10(8): p. e0135577, PMCID: PMC4532469, PMID: 26263555

40. Pasumarthi, K.B., E. Kardami, and P.A. Cattini, High and low molecular weight fibroblast growth factor-2 increase proliferation of neonatal rat cardiac myocytes but have differential effects on binucleation and nuclear morphology. Evidence for both paracrine and intracrine actions of fibroblast growth factor-2. Circ Res, 1996. 78(1): p. 126–36, PMCID: PMID: 8603495

41. Virag, J.A., M.L. Rolle, J. Reece, S. Hardouin, E.O. Feigl, and C.E. Murry, Fibroblast growth factor-2 regulates myocardial infarct repair: effects on cell proliferation, scar contraction, and ventricular function. Am J Pathol, 2007. 171(5): p. 1431–40, PMCID: PMC2043505, PMID: 17872976

42. Jensen, O.N., M. Wilm, A. Shevchenko, and M. Mann, Sample preparation methods for mass spectrometric peptide mapping directly from 2-DE gels. Methods Mol Biol, 1999. 112: p. 513–30, PMCID: PMID: 10027274

43. Tian, J., R. Wang, Q. Hou, M. Li, L. Chen, X. Deng, Z. Zhu, Y. Zhao, W. He, and X. Fu, Optimization and enrichment of induced cardiomyocytes derived from mouse fibroblasts by reprogramming with cardiac transcription factors. Mol Med Rep, 2018. 17(3): p. 3912–3920, PMCID: PMID: 29257325

44. Saladin, A., J. Rey, P. Thevenet, M. Zacharias, G. Moroy, and P. Tuffery, PEP-SiteFinder: a tool for the blind identification of peptide binding sites on protein surfaces. Nucleic Acids Res, 2014. 42(Web Server issue): p. W221–6, PMCID: PMC4086095, PMID: 24803671

45. Naito, H., I. Melnychenko, M. Didie, K. Schneiderbanger, P. Schubert, S. Rosenkranz, T. Eschenhagen, and W.-H. Zimmermann, Optimizing engineered ehart tissue for therapeutic applications as surrogate heart muscle. Circulation, 2006. 1114: p. I–72–I–78, PMCID: PMID:

