## Supplementary figures and images for "Identification and characterization of a Fibrillin-1 derived matrikine for cardiac regeneration and repair"

### Supplemental Figures 1 and 2

#### Supplemental Figures

#### Supplemental Figure 1

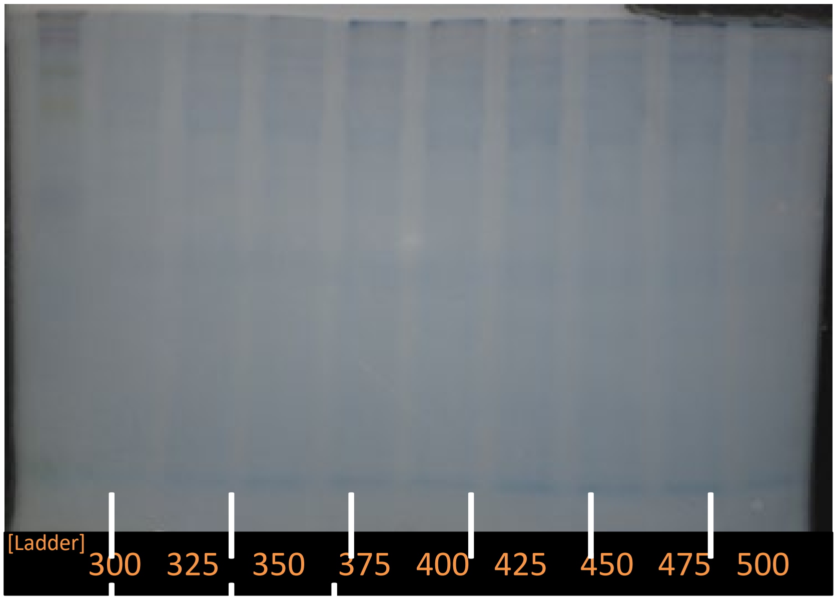

#### Supplemental Figure 2

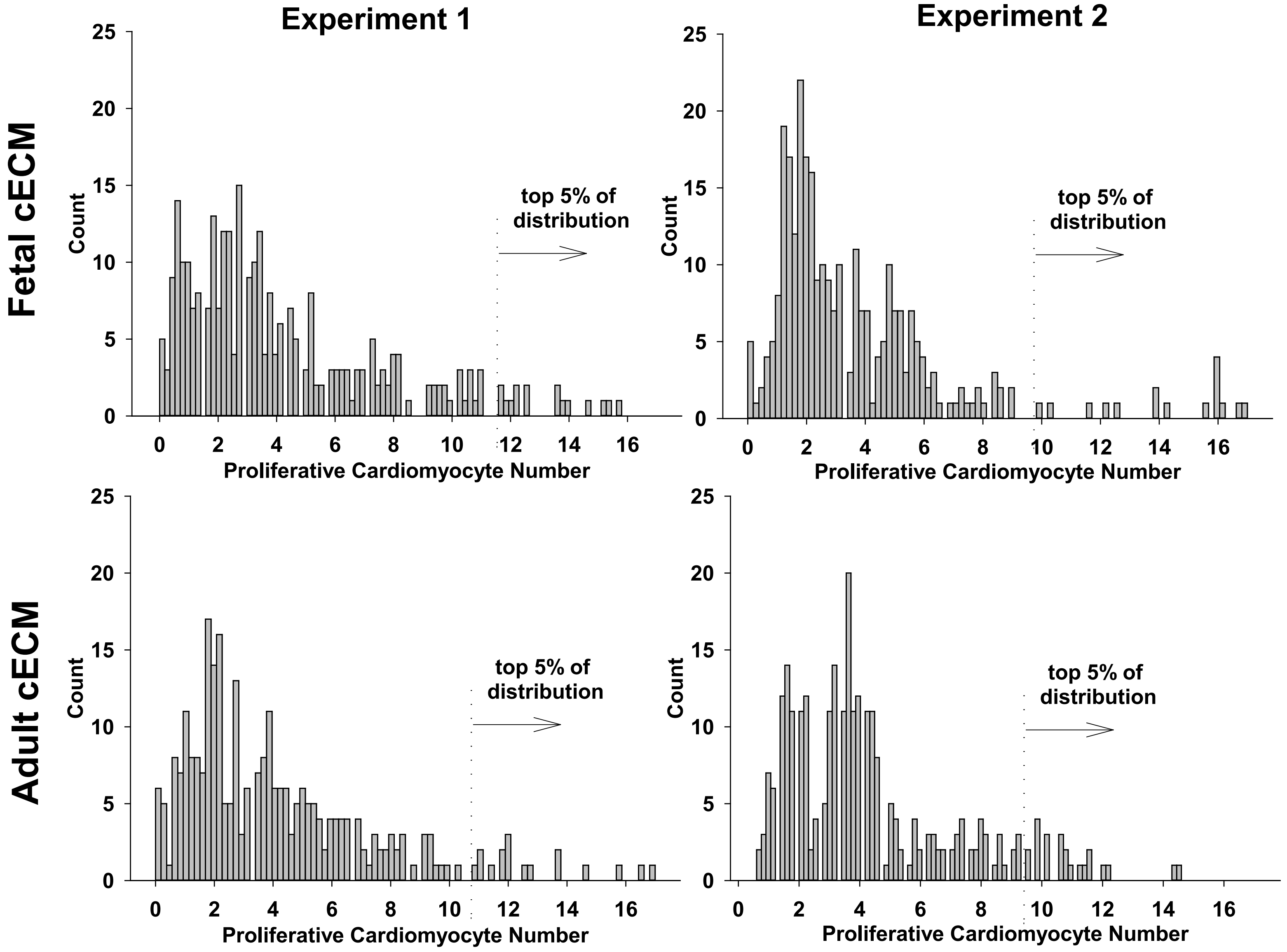
